## Supplementary Figures for "Tracking karyotype dynamics by flow cytometry reveals *de novo* chromosome duplications in laboratory cultures of *Macrostomum lignano*"

Suppl. Figure 1

A

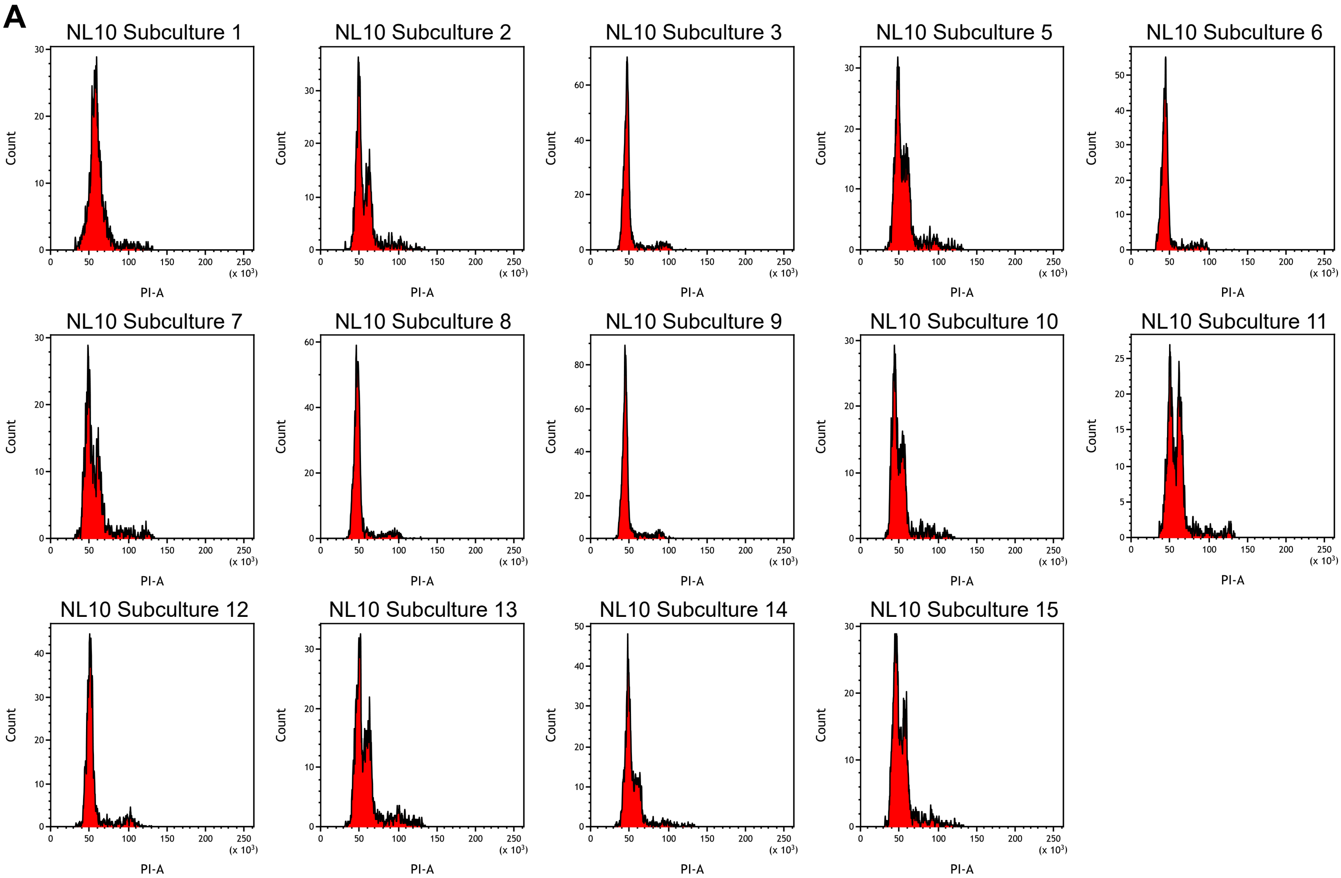


B

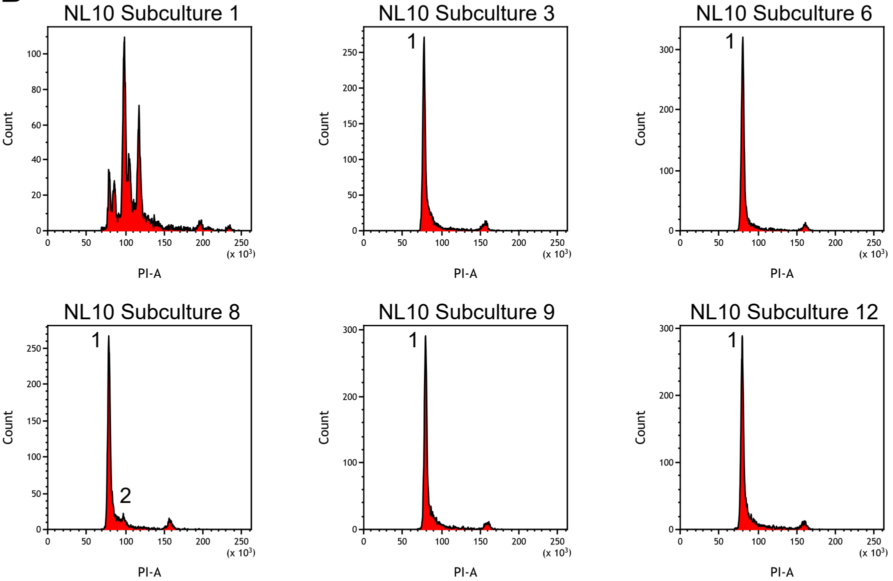

Suppl. Figure 2

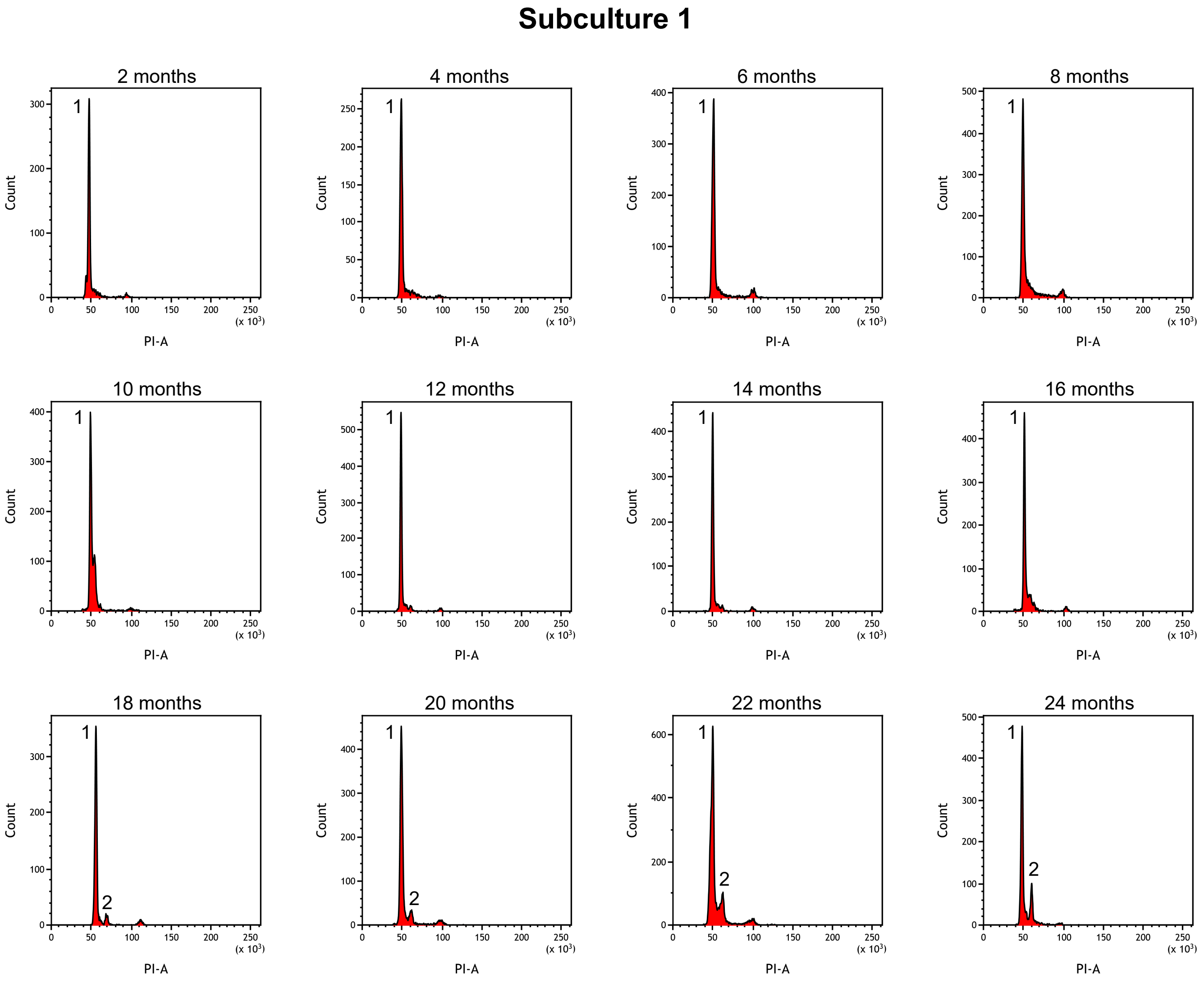

Suppl. Figure 3

#### Subculture 2

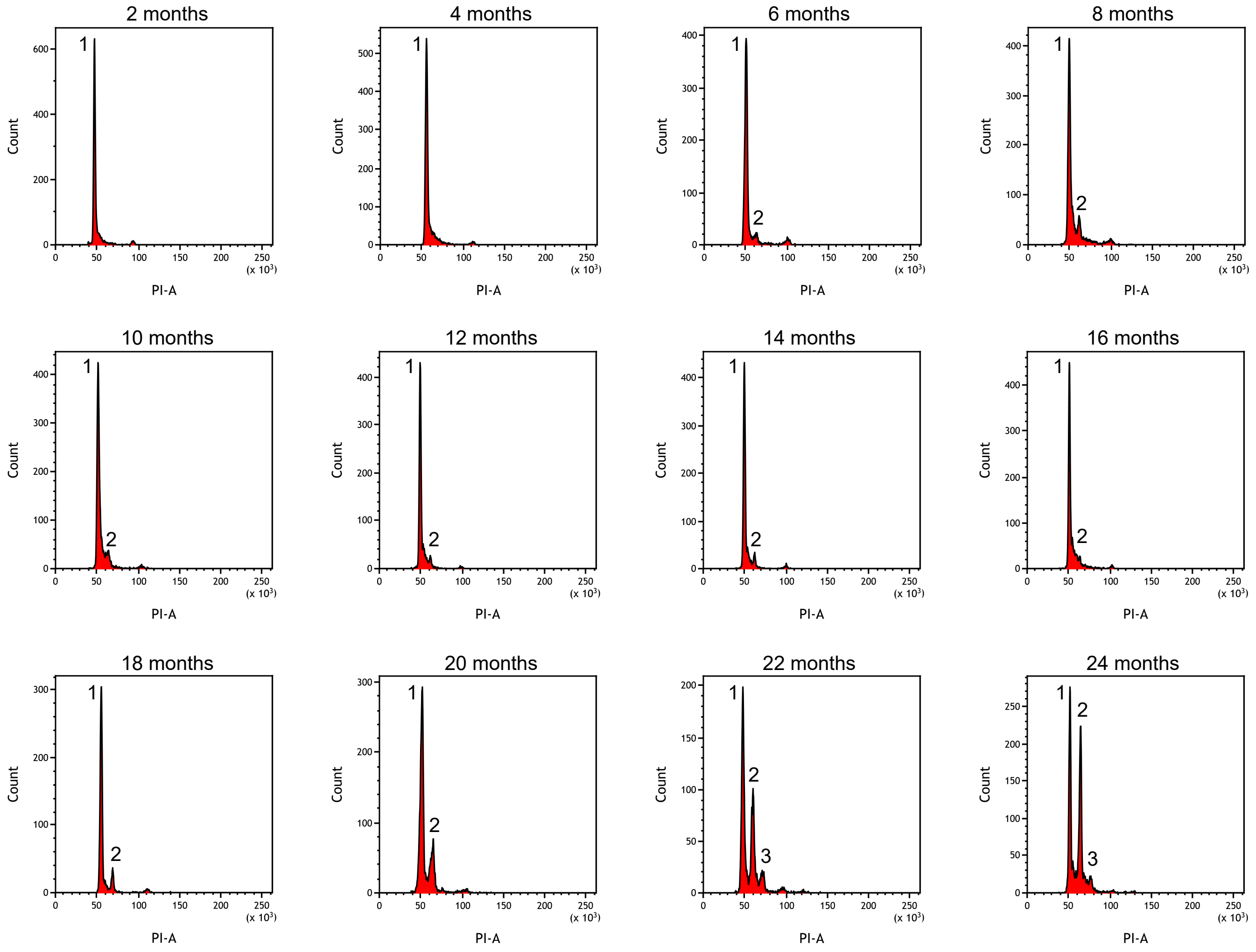

Suppl. Figure 4

#### Subculture 3

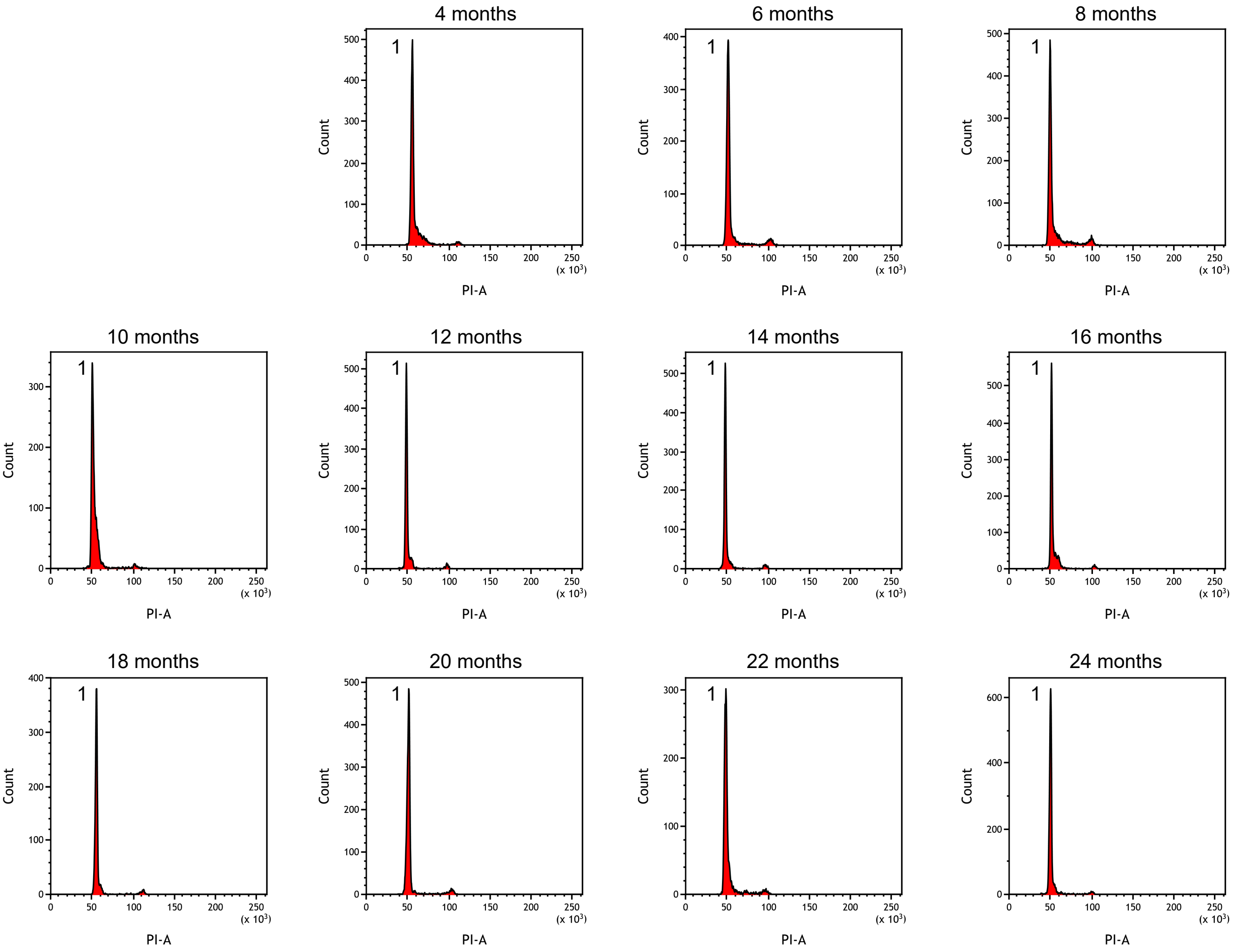

### Suppl. Figure 5

#### Subculture 4

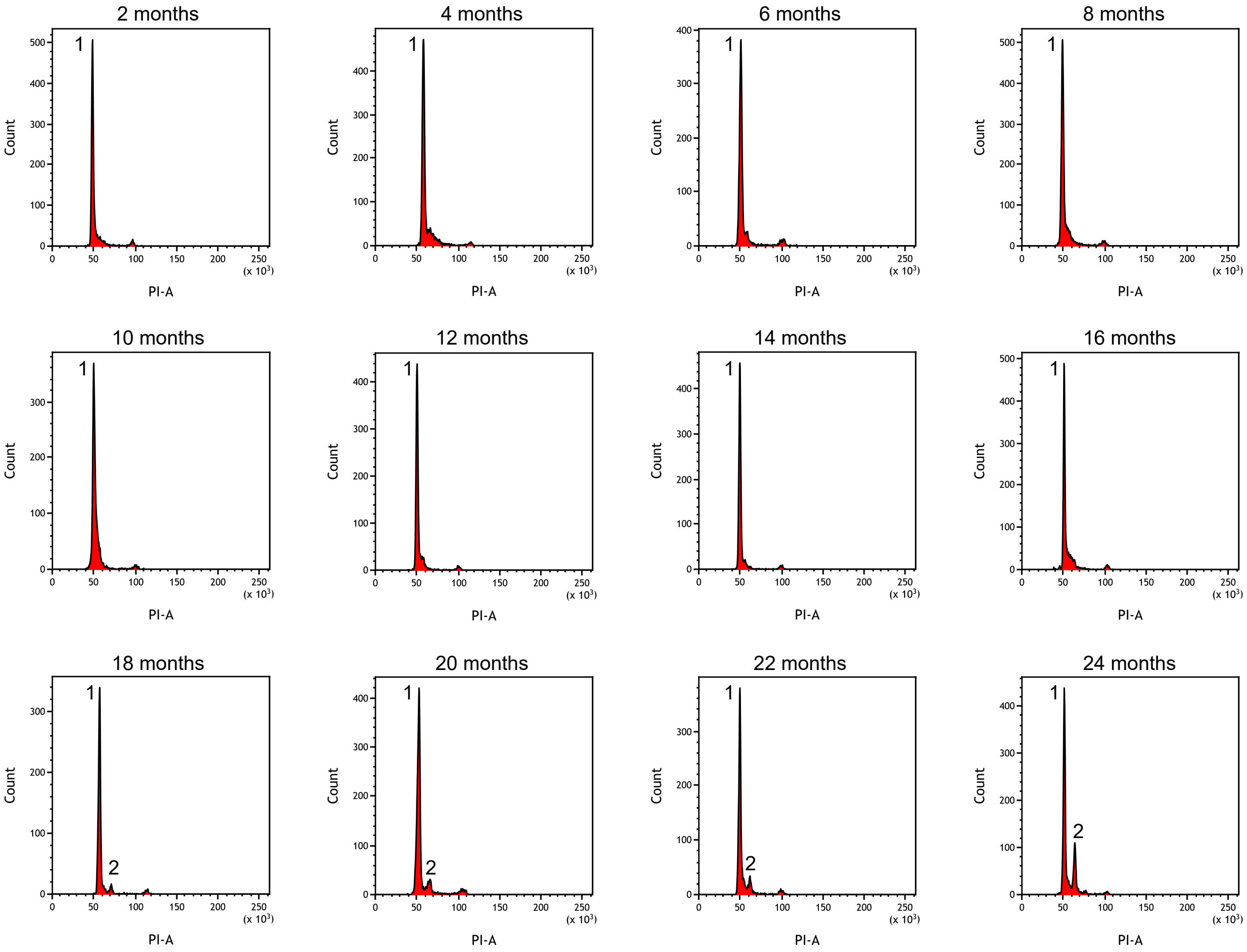

Suppl. Figure 6

#### Subculture 5

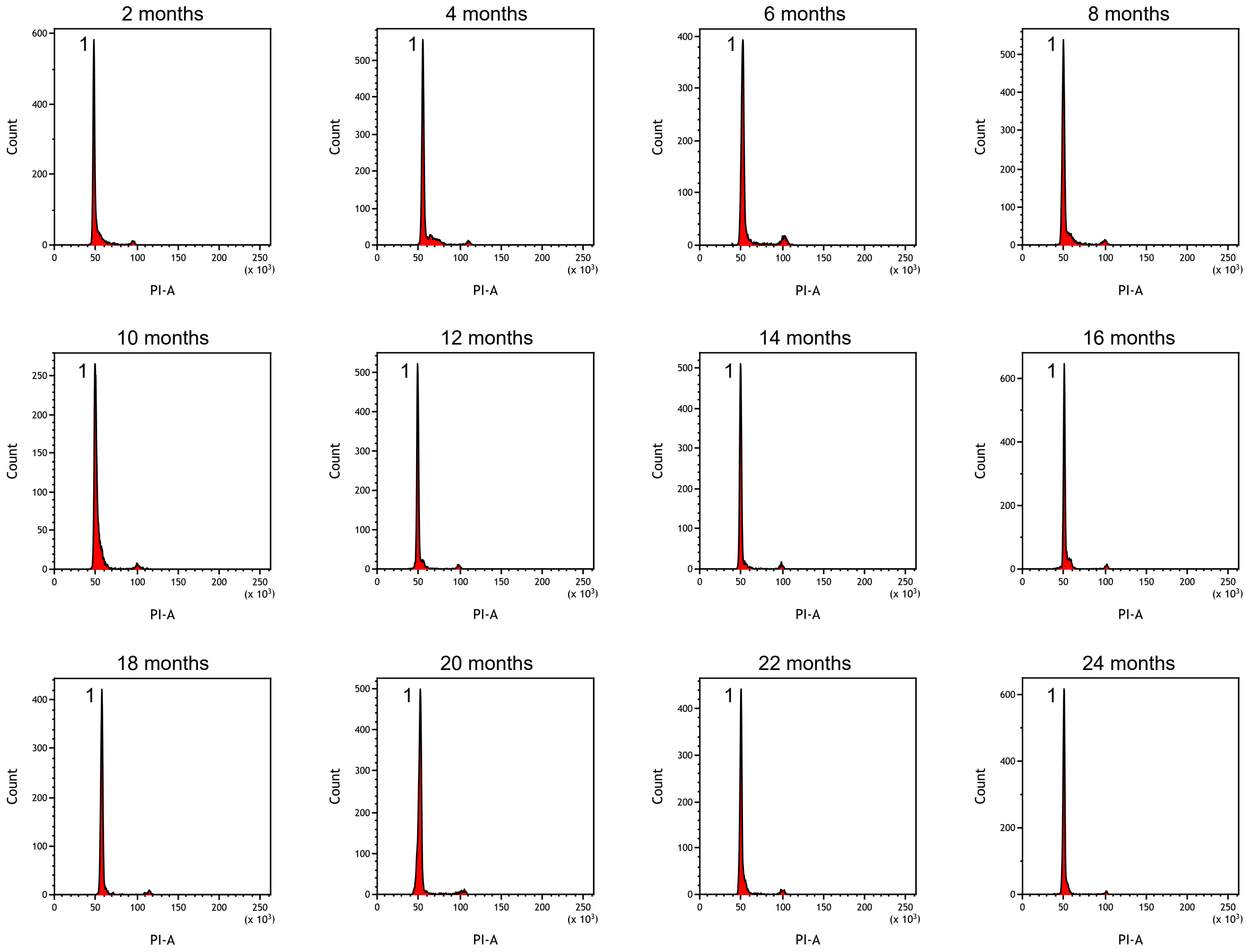

Suppl. Figure 7

#### Subculture 6

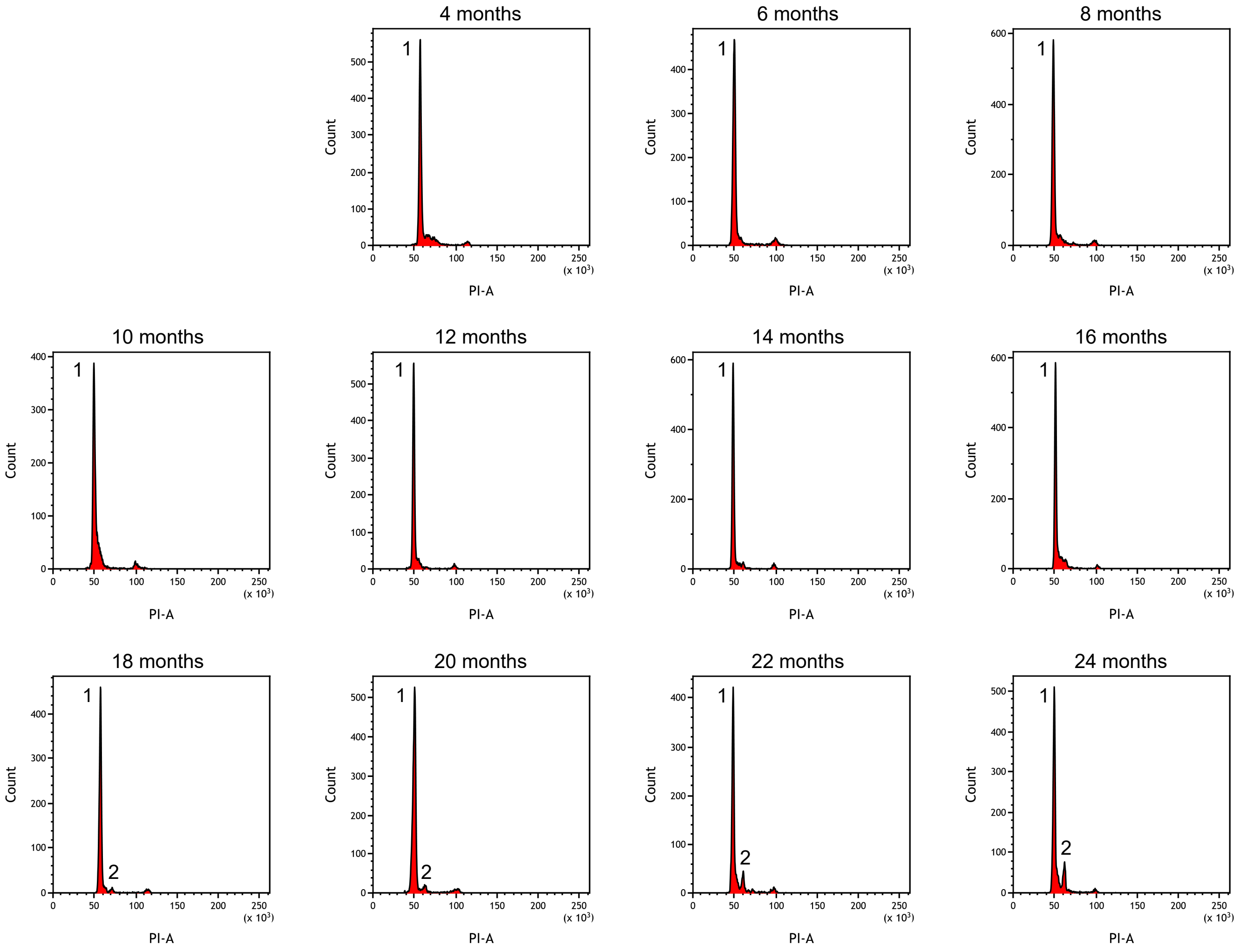

Suppl. Figure 8

#### Subculture 7

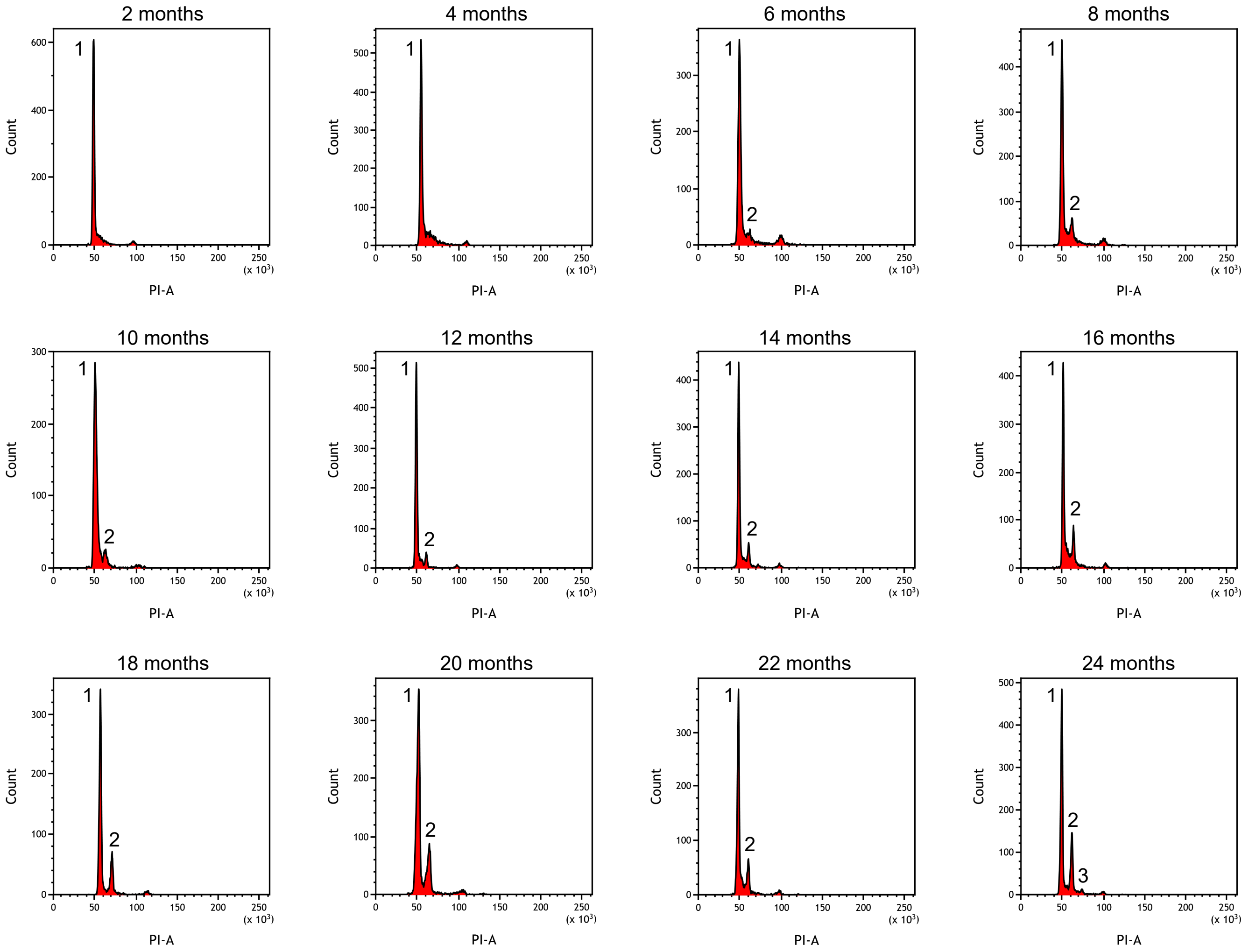

Suppl. figure 9

#### Subculture 8

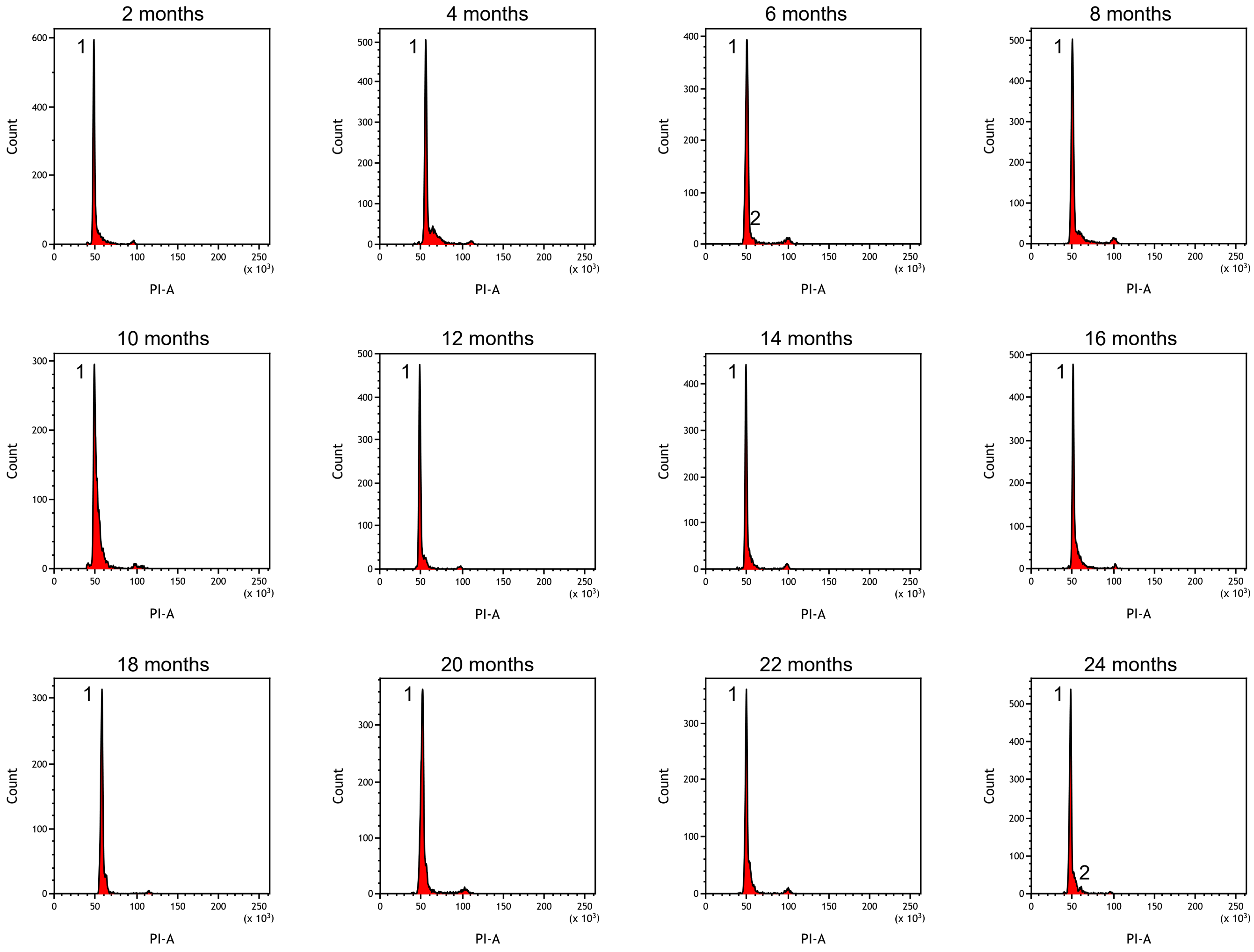

Suppl. Figure 10

#### Subculture 9

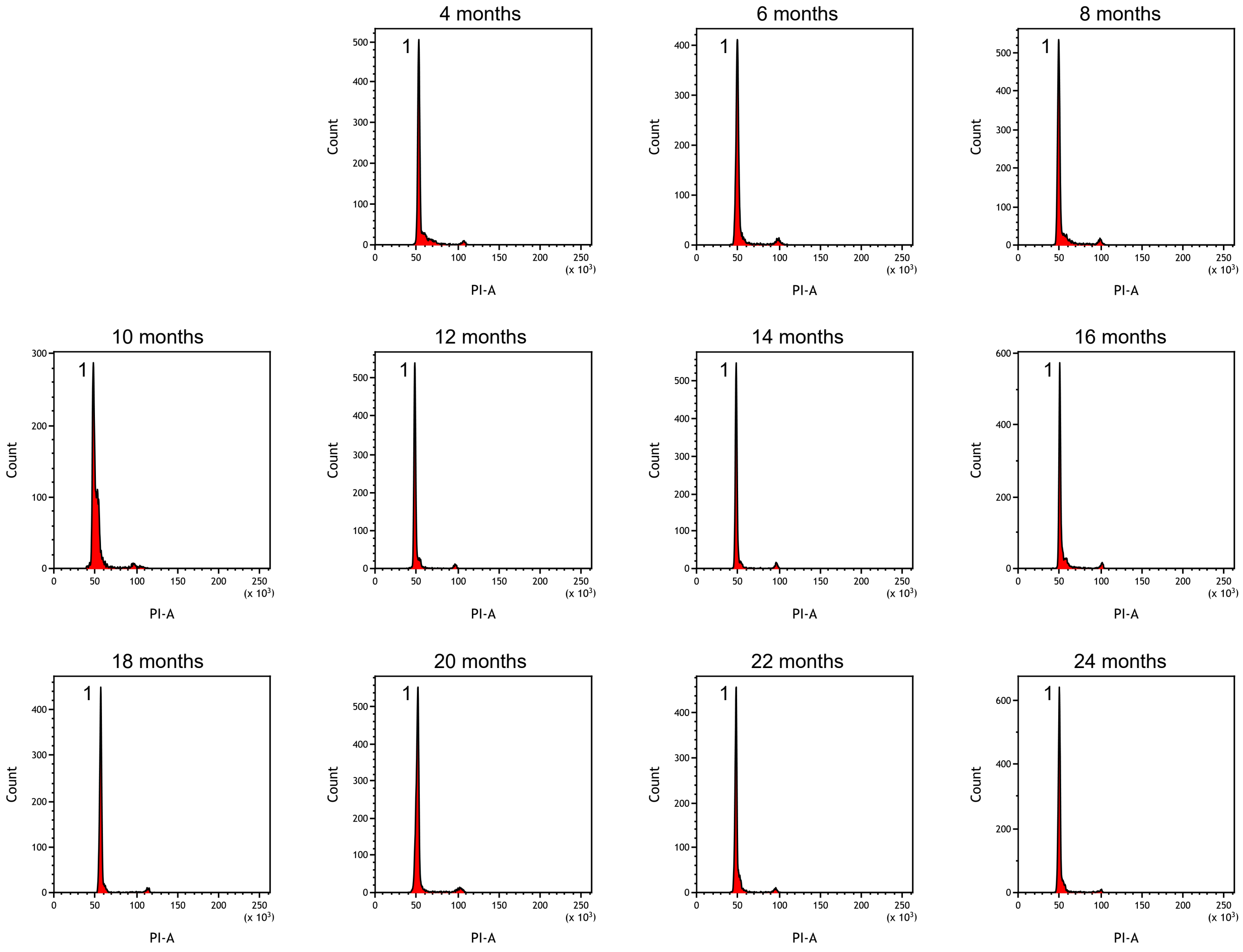

Suppl. Figure 11

#### Subculture 10

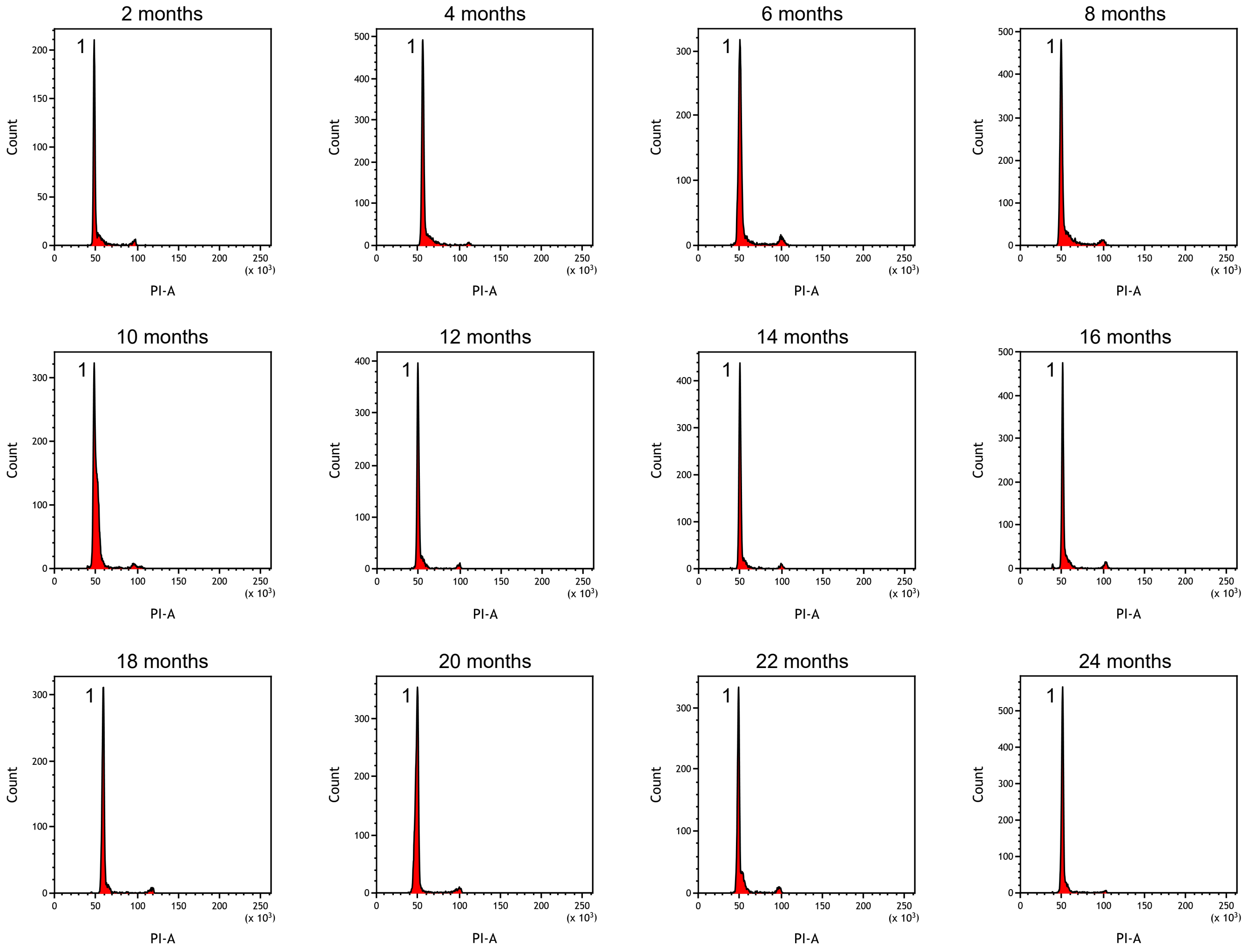

Suppl. Figure 12

#### Subculture 11

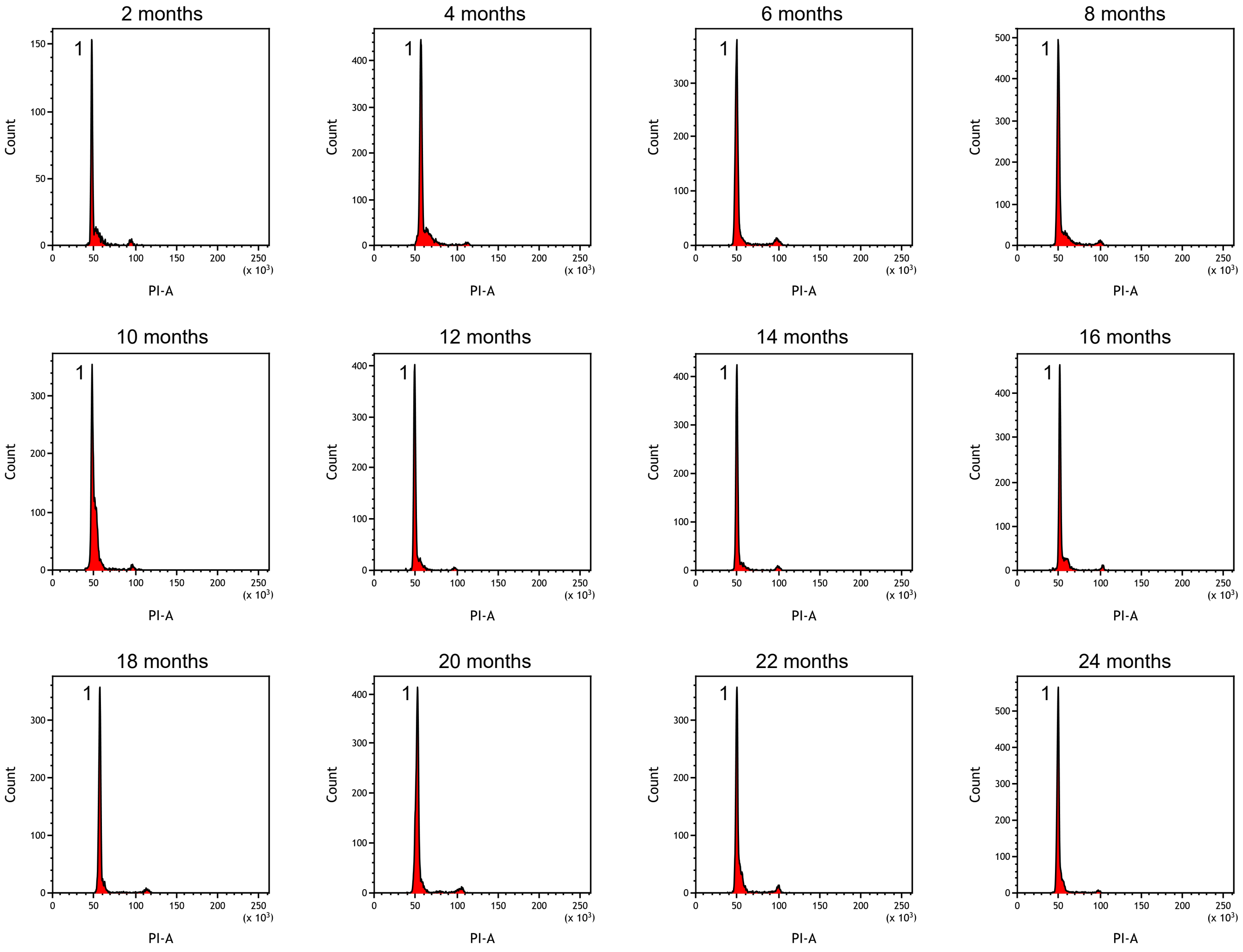

Suppl. Figure 13

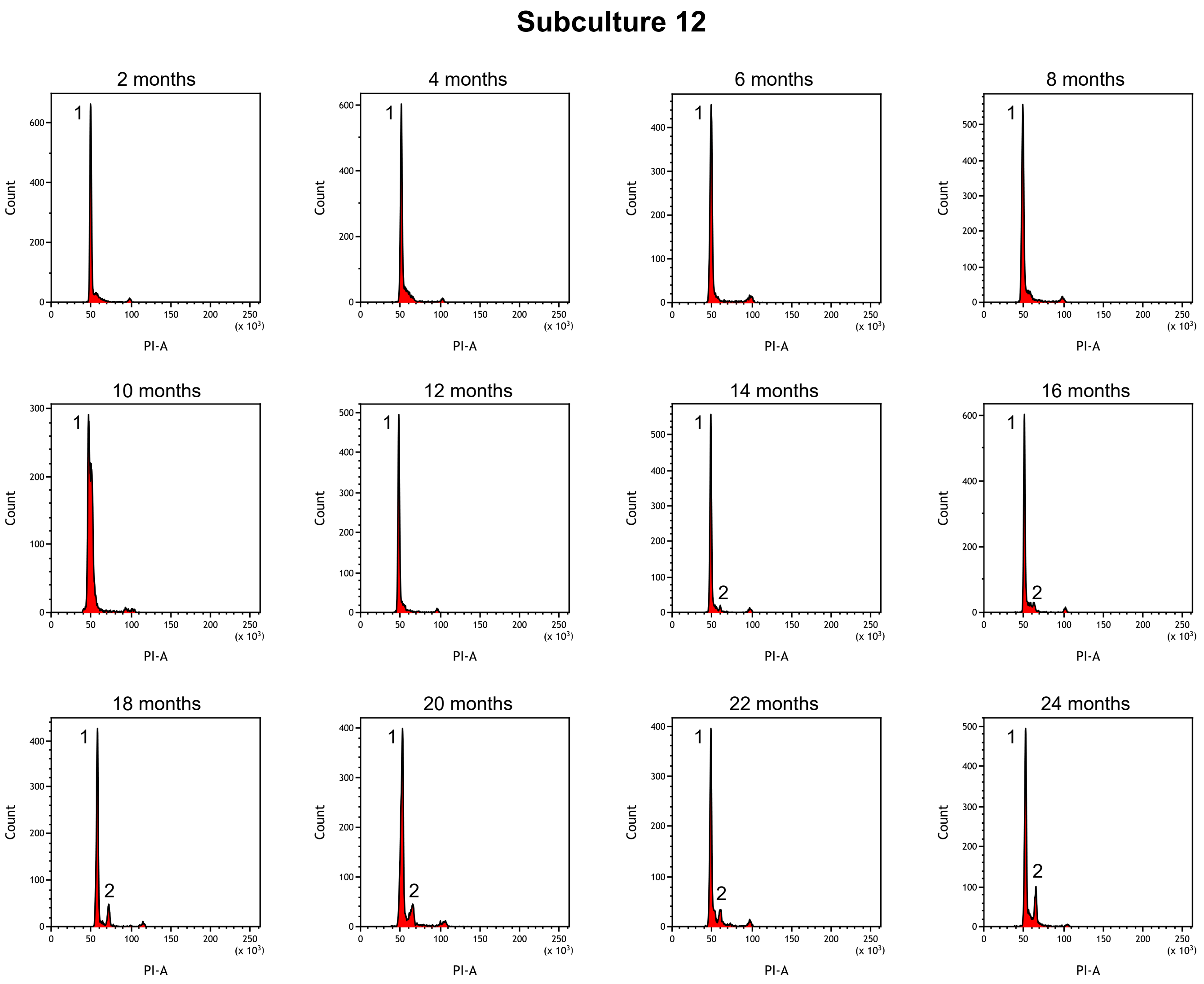

Suppl. Figure 14

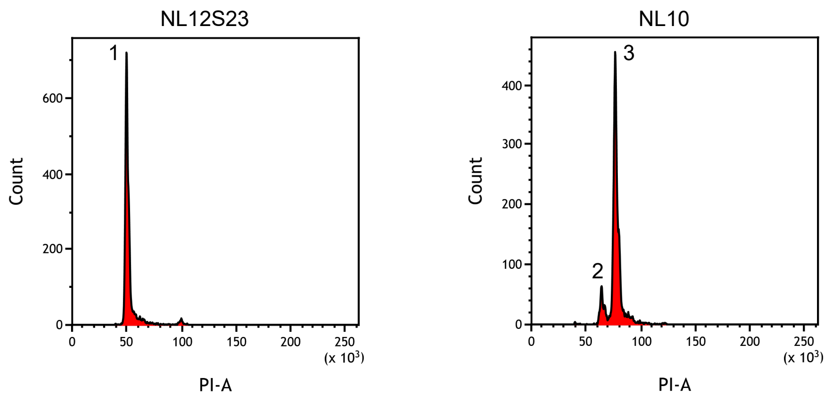
